# ProtTox: Toxin identification from Protein Sequences

**DOI:** 10.1101/2020.04.18.048439

**Authors:** Sathappan Muthiah, Debanjan Datta, Mohammad Raihanul Islam, Patrick Butler, Andrew Warren, Naren Ramakrishnan

**Author notes:** Equal Contribution (Sathappan Muthiah, Debanjan Datta).

## Abstract

Toxin classification of protein sequences is a challenging task with real world applications in healthcare and synthetic biology. Due to an ever expanding database of proteins and the inordinate cost of manual annotation, automated machine learning based approaches are crucial. Approaches need to overcome challenges of homology, multi-functionality, and structural diversity among proteins in this task. We propose a novel deep learning based method ProtTox, that aims to address some of the shortcomings of previous approaches in classifying proteins as toxins or not. Our method achieves a performance of 0.812 F1-score which is about 5% higher than the closest performing baseline.

## Introduction

Protein sequences, consisting of amino acids, are responsible for a multitude of biological processes such as metabolic reactions within cells and overall functioning at organism level. Proteins interact with other proteins, carbohydrates, lipids and DNA to regulate and facilitate various physiological and metabolic processes such as substrate channeling and enzyme activation. While many are responsible for maintaining the functional aspects of respective organisms, some have characteristics that render them toxic. *Toxins* are substances such as enzymatic or non-enzymatic proteins, which are deleterious to certain target organisms, produced through either biological processes in animals, plants and microorganisms or engineered through biotechnology. The task of classifying proteins as toxins or non-toxins is crucial to understanding which proteins are potentially hazardous. Further, knowing which protein sequences are toxins is advantageous in applications such as synthetic biology and healthcare.

With the advent of modern high throughput sequencing, the cost of screening many types of samples, clinical; environmental; experimental, is no longer prohibitive. As a result sequences are being made available to researchers in ever greater quantity. Since manual protein function annotation based on biological experiments is expensive and time consuming, the daunting challenge of annotating protein sequences and understanding their biological function still persists.

Machine learning based approaches can help address the challenges and provide an effective solution for the task of classifying protein sequences as toxins and help discover unknown features important for their identification. Machine learning models can be provided engineered features that encapsulate precious domain knowledge or they can implicitly accomplish feature extraction through representation learning. The latter is especially true for deep learning based models, which can efficiently perform feature extraction without the need for expensive feature engineering [2].

In our proposed model ProtTox, we adopt a supervised approach for the task of toxin identification in order to leverage the existing database of annotated protein sequences.

## Related Works

There have been multiple efforts in the past to overcome the limitations posed by experimental approaches by employing computational methods for determining protein function. It has been observed that proteins which share structurally similar regions (homology) demonstrate similar functionality. Homology or motif discovery based based clustering has been performed to group proteins which utilizes this concept, for instance *UniRef* clusters in *UniProt* database. Such methods have had limited success [9] in toxin prediction, due to the varying nature of proteins and structural similarity to non toxins owing to factors such as shared evolutionary lineage [8].

There have been specific machine learning based approaches proposed for toxin prediction. Clan-Tox[11] uses an ensemble of 10 classifiers with boosting to predict toxins. Global features are extracted from the sequence data and are used as inputs to a meta-classifier. ToxClassifier [7] also uses an ensemble based approach. It uses multiple approaches to engineer feature extraction which include techniques such as amino acid pair frequency, *BLAST* [10] and *HMMER* [15] based similarity scores.

Toxify [6] is a deep learning based approach for toxin prediction. It uses Gated Recurrent Networks, which have been effective for sequence classification in the natural language processing domain. Feature engineering here consists of obtaining Atchley Factors, that represents characteristics of each amino acid such as polarity and hydrophobicity. The authors demonstrate state of the art performance on their benchmark datasets indicating deep learning approaches can be an effective tool for our task. Our proposed model also employs a deep learning approach and we describe the details in the next section.

## Method

In this section, we describe our methodology for toxin prediction. Specifically, given a dataset *D* =< *s*_1_, *s*_2_, …, *s_n_* > of amino acid sequences, the task is to predict if any of the sequences, *s_k_*, is a toxin (the positive class) i.e., estimate *P*(*y_i_* = +1|*s_k_*, *θ*) where *θ* represents the model parameters. Figure. 1 provides an overview of three major components of our framework —(i) language model (*LM*), (ii) BiLSTM encoder (*E*) and finally, (iii) aggregation (*A*) and classification layer (*C*).

**Figure 1:**
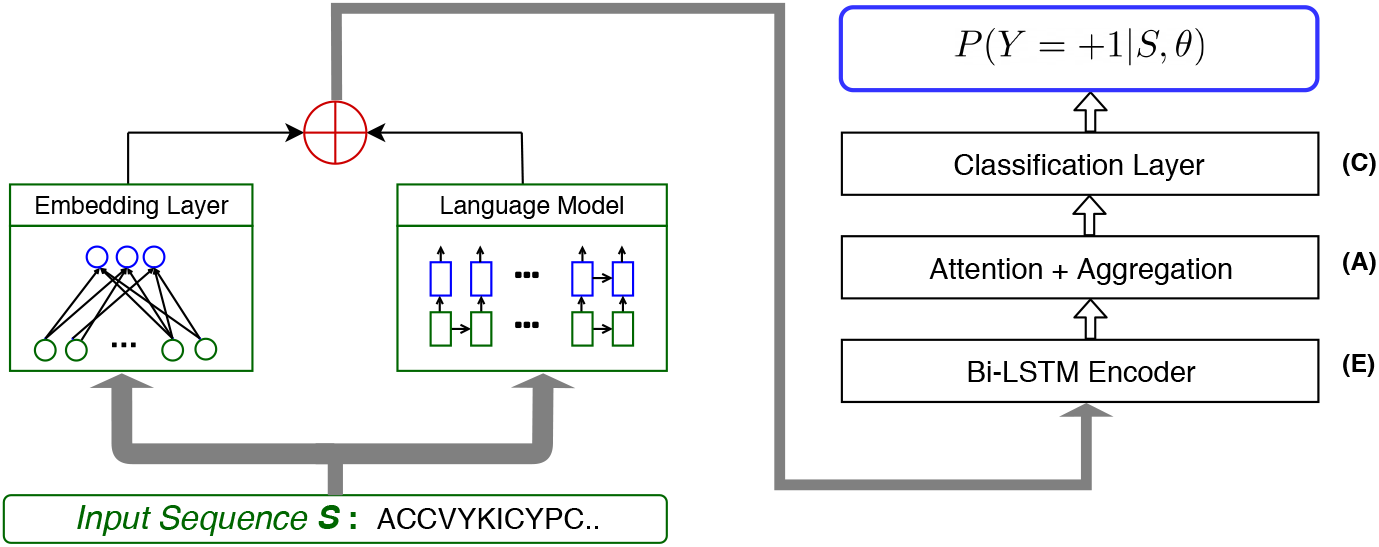
ProtTox Model Architecture

We make use of the pre-trained model from Tristan et al. [3] for the language model *LM* and the encoder *E*. The language model *LM* comprises of a bidirectional LSTM unit and was trained with the goal to predict the amino acid at a position *i* given information about the amino acids occurring before and after position *i* in the sequence. For our purpose we keep the language *LM* fixed and do not update its weights during training. The encoder unit *E* also comprises of Bi-directional LSTM units. *E* is fed as input a vector *h_i_* at each position *i*. The vector *h_i_* is a non-linear Rectified Linear Unit (ReLU) transformation of a combination of the language model output 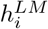 and an embedding vector *x_i_* for the amino acid occurring at position *i*.

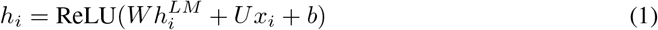

Next, for obtaining *P*(*Y* = + 1|*h_i_*), we first perform a linear transform of *h_i_* followed by SELU non-linearity and then obtain the mean along the length of the sequence to obtain a representation for the entire sequence, *e_s_*. Finally to obtain the toxin score, we pass *e_s_* through a dense layer followed by a sigmoid non-linearity. We call this overall model ProtTox.

Performing mean reduction over *h_i_* assumes every part of the sequence is important for making the prediction. However this may not be true. One way to identify which parts of the sequence are important is to make use of the attention mechanism. Specifically, we use dot-product attention [13], with a sigmoid activation instead of softmax, over *h_i_* to obtain attention probabilities. Additionally, to introduce sparsity and smoothness in the attention we train this model with an additional fused LASSO term. The overall loss function is shown in Eq. 2. Here *L* refers to sequence length and *a_i_* the attention probability for position *i*. We call this updated model with attention — ProtTox-Attn.

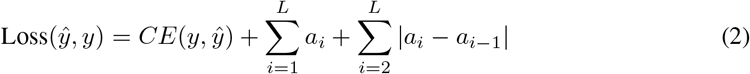

## Experimental Setup

### Data

All our models are trained on positive examples obtained from a combination of protein records with the toxin keyword “KW-0800” from Swissprot [4], and records marked as toxins from T3DB [16], and VFDB [5]. For the negative examples we used the the dataset from ToxClassifier [7] comprised of non-toxic sequences also from Swissprot. Specifically the ToxClassifer based negative dataset comprises of sequences drawn randomly from Swissprot (ignoring sequences that match the query *keyword: toxin OR annotation:*(*type: “tissue specificity” venom*)). The dataset also includes hard negative examples (with high similarity to toxin-like proteins but with physiological functions) obtained via a BLASTp search with *e*-value cutoff of 1.0*e* – 10.

For testing, we obtain a random held out sample of 1000 sequences with the Uniprot keyword “KW-0800” as the positive set and another 1000 sequences from PATRIC’s [14] metabolically essential genes list as the negative set.

### Methods for Comparison

We compare our method against the following state-of-art methods:

- **ToxClassifier** [7] is an ensemble of SVM and GBM with multiple engineered features.
- **Toxify** [6] is a deep recurrent model built over protein sequences where each amino acid is represented by the five Atchley factors [1]. Toxify requires the maximum sequence length to be 500, as such for when running Toxify we truncate large sequences at position 500.
- **SWEM** [12] is a simple word embedding model that has been shown to be effective in text classification approaches. We adapt the SWEM model for toxin classification. Size 3 k-mers are used as input to the SWEM model.
- **ProtTox** and **ProtTox-Attn** are our methods described in previous section.

Finally, we train all models with 3-fold cross-validation.

## Results

In this section, we present the results of our experimental analysis. Specifically we would like to answer the following questions:

### 1) What is the performance of ProtTox on toxin prediction task?

Table. 1 shows the performance of the different models in our test set. We observe that ProtTox-achieves the best precision scores for toxin prediction, whereas ProtTox-Attn achieves a near-perfect score of 99.8%. We notice that both models have lower recall score compared to SWEM, which slightly hurts the F1 scores for both models. However, for non-toxin prediction ProtTox-Attn achieves the best F1 score. This demonstrates our proposed models can be fruitful for vanilla toxin prediction.

**Table 1:** The *ProtTox* approach achieves the best precision and F-1 score for Non-Toxin prediction. For toxin prediction, our methods achieves comparable performance w.r.t to the SWEM model

| ModelNames | Non-Toxin |  |  | Toxin |  |  |
| --- | --- | --- | --- | --- | --- | --- |
|  | Precision | Recall | F-1 | Precision | Recall | F-1 |
| ToxClassifier | 0.620 | **1.000** | 0.760 | **1.000** | 0.390 | 0.560 |
| Toxify | 0.530 | 0.860 | 0.660 | 0.790 | 0.410 | 0.540 |
| SWEM | **0.723** | 0.957 | 0.824 | 0.936 | **0.634** | **0.756** |
| ProtTox | 0.686 | **1.000** | 0.814 | **1.00** | 0.542 | 0.703 |
| ProtTox-Attn | 0.706 | 0.999 | **0.827** | 0.998 | 0.584 | 0.737 |

### 2) Does predicting sub-toxin keywords help improve overall Toxin Prediction results?

In this experiment, we try to induce more structure into the prediction task by breaking down the Toxin class into subtypes. For this we make use of the Uniprot keyword hierarchy to identify 11 sub-types: Cardiotoxin, Cell adhesion impairing toxin, Complement system impairing toxin, Dermonecrotic toxin, Enterotoxin, G-protein coupled receptor impairing toxin, Hemostasis impairing toxin, Ion channel impairing toxin, Myotoxin, Neurotoxin and finally for any sequence that is a toxin but does not fall into any of the above mentioned subtypes were put under a class called “Other”.

For this task, we only compare the performance of SWEM and ProtTox-Attn(the best performing models from Table. 1). The models are trained with cross entropy loss where the overall score for toxin is obtained by summing up the individual sub-type scores. Figure. 2(A) shows the performance on toxin identification when we also try to predict the sub-type. We notice that the introduction of additional structure to the prediction problem (in terms of sub-types and their hierarchical relation with the root toxin class) helps ProtTox-Attn to learn better representations for toxin classification. We believe adding sub-type predictions have helped regularize and learn a better minima for our model ProtTox-Attn. However we note that the SWEM model performs poorly under this setting. This is possibly due to the fact the SWEM model, being single layered model, does not have enough complexity/capacity to model this increased label space.

**Figure 2:**
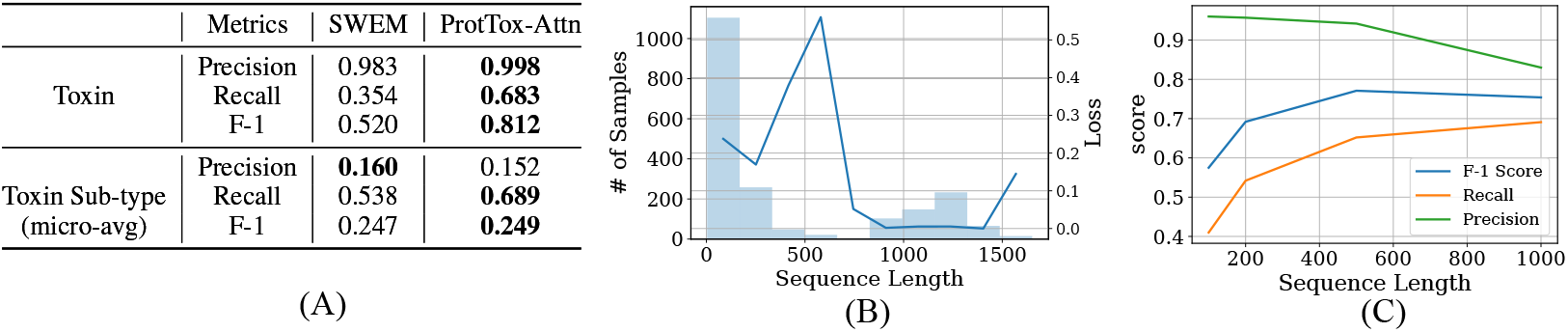
(A) provides the performance metric comparison between SWEM and ProtTox-Attn when the models are trained for sub-type prediction also. With sub-type prediction the overall toxin prediction capability of ProtTox-Attn improves. We see the best f1-score of **0.812** which is 5% better than the best SWEM model (from Table. 1). (B) shows how ProtTox-Attn performs w.r.t to sequences of different length. The bar indicates the # of samples with a given sequence length, whereas the curve indicates the average loss incurred for sequences with a given length. (C) shows the performance difference when SWEM model is trained with different sequence length cut-offs.

### 3) What is the effect of sequence length on toxin prediction?

In order to answer this question, we perform two sets of experiments. First, we train multiple SWEM models (chosen for its simplicity) with max sequence length set to 100, 200, 500, 1000. Under each setting, if a sequence has a length greater than the specified length then its truncated. The results of this experiment are shown in Figure. 2(C). We notice that with increasing sequence length precision decreases but recall increases. As such, we observe optimal F1-score somewhere in the middle around sequence length 500. Secondly, on Figure. 2(B) we investigate how the loss of ProtTox-Attn varies w.r.t to sequence length. We notice that our model performs best for long sequences.

## Conclusion

In this paper, we develop deep a learning based approach for toxin prediction. We observed that predicting toxin sub-types help improve overall toxin classification performance especially with increasing complexity (depth) of the model. We also study the performance variation of our model in terms of sequences of different lengths. Moving forward, it is possible to extend our model in several directions. Incorporating richer domain knowledge, such as on PPI and 3-D structure of proteins etc., can potentially provide greater predictive power.

## Acknowledgements

This study is based upon work supported by the Office of the Director of National Intelligence (ODNI), Intelligence Advanced Research Activity (IARPA), via the Army Research Office (ARO) under cooperative agreement number W911NF-17-2-0105A.

